# In situ cryo-electron tomography of vaccinia virus exit from infected cells

**DOI:** 10.1101/2025.08.20.671218

**Authors:** Miguel Hernandez-Gonzalez, Andrea Nans, Thomas Calcraft, Peter Rosenthal, Michael Way

**Affiliations:** Cellular Signalling and Cytoskeletal Function Laboratory, The Francis Crick Institute, 1 Midland Road, London, NW1 1AT, UK; Structural Biology Science Technology Platform, The Francis Crick Institute.; Structural Biology of Cells and Viruses Laboratory, The Francis Crick Institute.; Department of Infectious Disease, Imperial College, London SW7 2AZ, UK

## Abstract

Poxvirus-infected cells release newly assembled virions via Golgi-mediated envelopment and subsequent exocytosis at the plasma membrane, prior to cell lysis. During exocytosis, the viral outer membrane fuses with the plasma membrane but retains a distinct identity, beneath which septins and clathrin are independently recruited. Clathrin enhances actin-dependent viral spread, while septins suppress virus release from the cell. Here, we used cryo-electron tomography and structured illumination microscopy (SIM) to study vaccinia egress. Our 3D analysis reveals that Golgi-mediated envelopment is a highly adaptable process that involves remodelling of the enfolding membrane and a final pore sealing step. Filaments that are ∼ 4-nm wide, which are likely septins based on their structural features and our SIM imaging, run parallel to the inner surface of the plasma membrane beneath virions attached to the cell surface. In contrast, clathrin induces plasma membrane invaginations at distinct subdomains beneath the virus. We propose that actin assembly at these clathrin subdomains would provide a template for subsequent virus-induced actin polymerization, therefore enhancing it.

## Introduction

Vaccinia is the prototypical member of the poxvirus family, which includes the causative agents of smallpox and mpox (Beiras et al., 2025; Gubser et al., 2004; Makokha et al., 2025). Poxviruses are enveloped double stranded DNA viruses that undergo replication and virion assembly in the cytoplasm of infected cells (Moss, 2007; Roberts and Smith, 2008). During the initial spread of infection, vaccinia virus leaves the cell without causing host cell lysis. To achieve this, newly assembled intracellular mature virions (IMV) get enveloped with a modified Golgi/endosomal compartment to form intracellular enveloped virions (IEV), consisting of a nucleoprotein core enveloped by three membranes (Schmelz et al., 1994; Smith et al., 2002; Tooze et al., 1993). The viral proteins required for IMV envelopment have been identified (Blasco and Moss, 1991; Dodding et al., 2009; Domi et al., 2008; Engelstad and Smith, 1993; Rottger et al., 1999; Smith et al., 2002; Wolffe et al., 1993), but we lack a structural understanding of the wrapping process that produces IEV.

Once formed, IEV are transported on microtubules by kinesin-1 to the plasma membrane to be released out of the cell by the fusion of the IEV outermost membrane with the plasma membrane (Geada et al., 2001; Hollinshead et al., 2001; Newsome et al., 2004; Rietdorf et al., 2001; van Eijl et al., 2002; Ward and Moss, 2001; Ward and Moss, 2004; Xu et al., 2023). The outermost IEV membrane, containing viral membrane proteins such as A36 which recruits kinesin-1 (Dodding et al., 2011; Rietdorf et al., 2001; Ward and Moss, 2004), becomes part of the plasma membrane but retains a distinct viral identity beneath the released virion (Roberts and Smith, 2008; Smith et al., 2002). This process releases a double-membraned extracellular virus, which can remain attached to the outer cell surface as cell-associated enveloped virions (CEV) (Roberts and Smith, 2008; Smith et al., 2002). Subsequently, CEV can induce an outside-in signalling cascade that induces Src and Abl-mediated phosphorylation of the viral protein A36 (Frischknecht et al., 1999; Newsome et al., 2004; Newsome et al., 2006; Reeves et al., 2005). This phosphorylation promotes dissociation of kinesin-1 and activates Arp2/3-driven actin polymerisation beneath the virion to enhance the cell-to-cell spread of infection (Cudmore et al., 1995; Doceul et al., 2010; Frischknecht et al., 1999; Moreau et al., 2000; Newsome et al., 2004).

Prior to the initiation of actin polymerisation, septins are recruited beneath CEV immediately after IEV fuse with the plasma membrane (Pfanzelter et al., 2018). Septins are an essential component of the cytoskeleton involved in many fundamental cellular processes including cytokinesis and cell migration (Cavini et al., 2021; Mostowy and Cossart, 2012; Schampera and Schwan, 2024; Spiliotis and McMurray, 2020; Woods and Gladfelter, 2021). They comprise a family of GTP binding proteins that form non-polar filaments, which can organise into higher order assemblies as well as interact with actin, microtubules and membranes. While high order structures that septins can form have been studied in vitro (Nakazawa et al., 2023), their organisation in cells, especially in association with actin, microtubules and membranes, remains unclear partly due to electron microscopy limitations. The molecular basis for septin recruitment beneath CEV is not established, but their presence suppresses the release of the virus from the surface of infected cells (Pfanzelter et al., 2018). In contrast, the ability of CEV to induce actin polymerisation is enhanced by the independent recruitment of clathrin after septins (Humphries et al., 2012; Pfanzelter et al., 2018). Clathrin and its plasma membrane binding adaptor AP-2 are recruited by Eps15 and intersectin-1, which interact with three NPF motifs in the cytoplasmic tail of A36 (Snetkov et al., 2016). It remains to be established how clathrin recruitment enhances the ability of CEV to induce actin polymerization. Clathrin typically forms polyhedral lattices on membranes (Traub, 2011; Vassilopoulos and Montagnac, 2024), whereas septin organisation at the plasma membrane remains unclear. However, confocal imaging of vaccinia-infected cells suggests septins form rings or cages beneath CEV, consistent with a supramolecular assembly (Pfanzelter et al., 2018). Understanding the molecular organisation of septins and clathrin beneath CEV will shed light on how these macromolecular assembles perform opposing functions during viral egress without mutual interference.

Here, we have studied the egress pathway of vaccinia virus using cryo-electron tomography (cryo-ET), as part of our broader interest in obtaining a better understanding of its replication cycle in infected cells (Hernandez-Gonzalez et al., 2023). We have obtained new insights into the envelopment process that generates IEVs and uncovered filaments which we believe are septins running parallel to the plasma membrane beneath CEV, based on their width and difference in appearance from actin or intermediate filaments. In contrast, clathrin forms canonical endocytic pits that deform the plasma membrane, which we propose act as initiation sites to enhance actin assembly during viral spread.

## Results

### Cryo-electron tomography of IMV envelopment during IEV formation

To visualise the different stages of viral egress, we imaged the electron-transparent periphery of vaccinia-infected HeLa cells using cryo-ET at 8 hours post-infection. We were able to capture tomograms containing all forms of viral particles (IMV, IEV and CEV). Importantly, we were also able to image IMV undergoing envelopment during IEV formation (Figures 1 and S1). The association of membrane cisternae with the IMV surface, which is also a membrane, could be observed at different stages, ranging from a small portion of the virus to examples where the IMV is completely covered. It was also noticeable that the rims of the wrapping cisterna were often dilated (Figure S1). When tomograms of fully wrapped IMV were examined in 3D, we observed three examples where there was a single pore of approximately 7 nm in diameter at the final point of wrapping (Figure 1A-B and Movie 1). After close examination, one of these enveloping IMV presented multiple pores (Figure S1B and Movie 2). For the virus to be released from the infected cell, these small apertures must be sealed before the outer IEV membrane fuses with the plasma membrane (Figure 1C). Consistent with this, we were able to image fully closed IEV near the plasma membrane (Figures 1D and S2).

**Figure 1.**
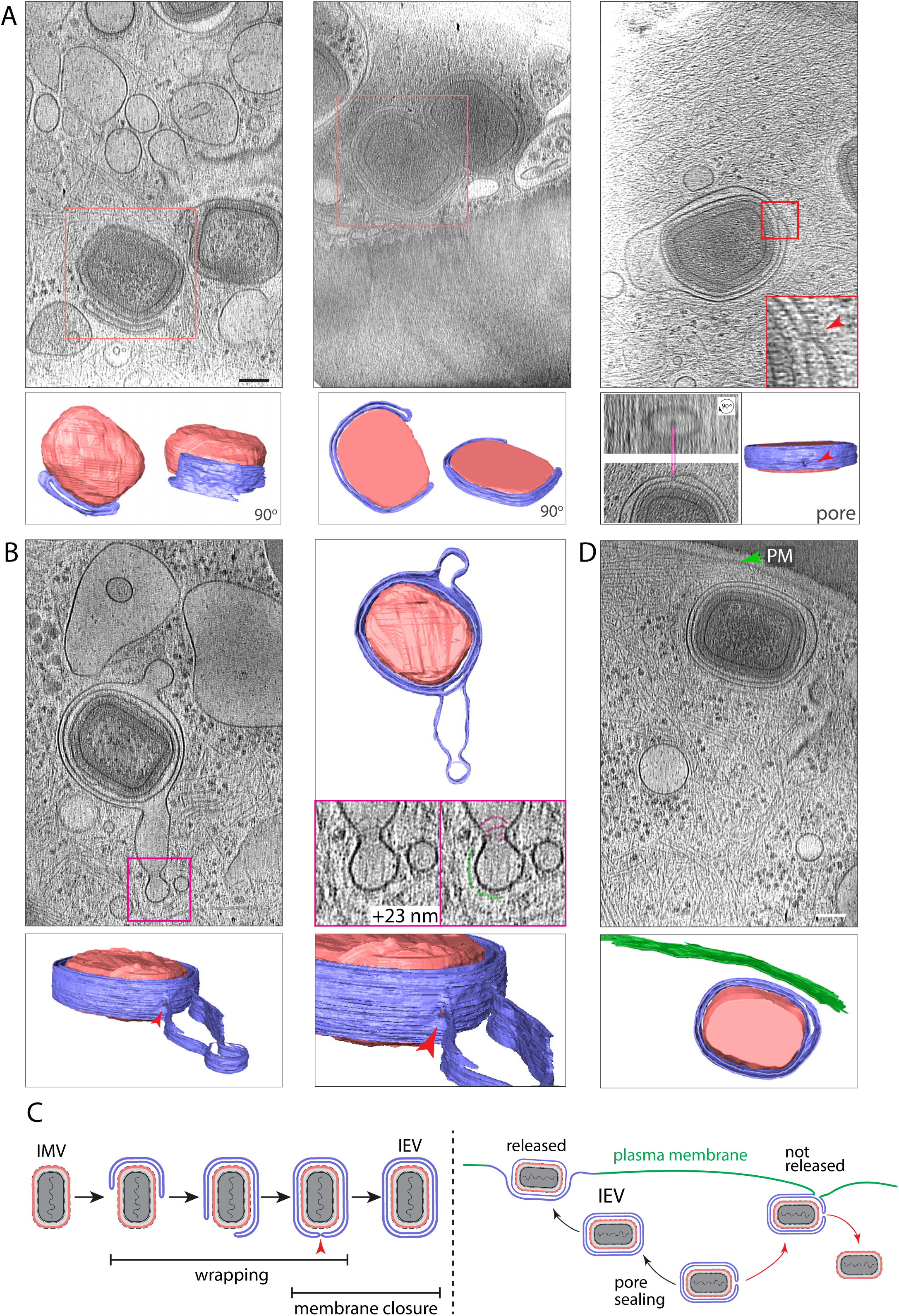
Cryo-ET of vaccinia virus envelopment. **A.** Middle sections of reconstructed tomograms showing different degrees of IMV envelopment inside vaccinia-infected cells. Red squares indicate the envelopment events segmented below. On the top right image, the red square corresponds to the zoomed in region in which a pore is present in the enveloping membrane (red arrowhead) (see Movie 1). Below, an orthogonal view of the same tomogram, focused on the pore, and its 90-degree rotated image. **B.** Mid-section of a tomogram displaying an envelopment event showing membrane tubulation and constriction. Collar-like structures and a putative coat are labelled in magenta and green, respectively. (See Movie 3). **C.** Schematic representation of vaccinia envelopment where a final pore is sealed by annular membrane fusion (left) and the consequences of not sealing the pore prior to exocytosis at the plasma membrane (right). While the rest of the model is based on previous work, here we propose a final pore occurs during this process, which must be sealed. **D.** Fully enveloped and sealed IEV, in close proximity to the plasma membrane indicated by the green arrowhead. A different section of the same tomogram has previously been shown in Figure 7 (Hernandez-Gonzalez et al., 2023). In all cases, 3D manual segmentations of tomograms are provided where pale red represents the IMV surface and the enveloping membrane cisternae is shown in blue. Red arrowheads indicate pores in the wrapping cisterna while the plasma membrane is shown in green. Scalebars = 100 nm.

A notable feature is the variability in the size of the membrane cisternae enveloping IMV. When excess wrapping membrane is present it does not prevent its inner surface from tightly associating with the IMV surface (Figures 1 and S1A-B). Moreover, we observed constrictions in the outer surface of the wrapping membrane suggesting that remodelling occurs to remove excess membrane during IMV envelopment (Figure 1B and Movie 3). Consistent with this, we were able to see collars of protein density at membrane constrictions. We also found constrictions of the outer membrane of fully sealed IEV (Figure S2B). In addition, at the plasma membrane, we found multiple IMV sharing a single envelope on the outside of the cell suggesting that envelopment is a flexible and adaptable process that can occur around multiple particles (Figure S2C). In agreement with this, additional material such as vesicles can get trapped inside the enveloped particle (Figures S1C and S2A).

### Cryo-electron tomography of CEV on the plasma membrane

To facilitate the analysis of CEV prior to actin polymerisation, we performed cryo-ET on cells infected with vaccinia A36-YdF strains that cannot induce actin polymerisation (Frischknecht et al., 1999; Ward and Moss, 2001). We obtained tomograms of top down and side views of CEV on the outside of infected cells (Figure 2A-C and Movies 4 and 5). Quantification of the percentage of the CEV surface in contact with the plasma membrane in the median plane of our tomograms reveals that the extent of invagination is variable (Figure 2D). In contrast, the distance between the CEV surface and the outer IEV membrane that has been incorporated into the plasma membrane is relatively constant at 11.52 ± 0.23 nm (Figure 2D). The numerous densities visible on the CEV and the underlying plasma membrane presumably represent the luminal tails of IEV proteins such as B5, which are responsible for retaining the CEV on the outer surface of the cell (Figure 2E).

**Figure 2.**
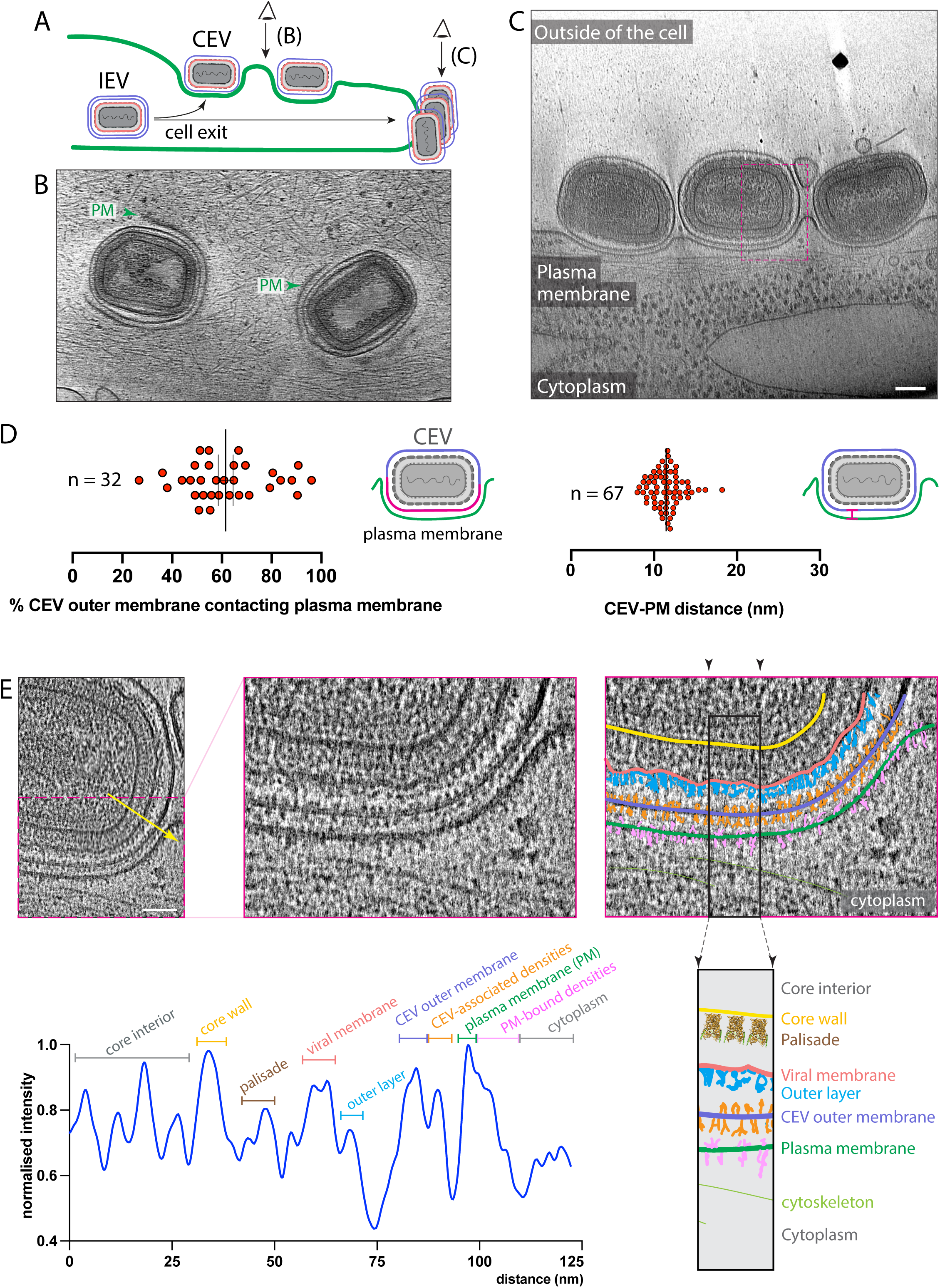
Cryo-ET analysis of cell-associated extracellular vaccinia virus (CEV). **A.** Schematic of IEV release from the infected cell. After exocytosis, the resulting CEV can be visualised by cryo-ET on top of the cell or at the cell edge, offering two different views exemplified by B and C. **B.** Top view of CEV on the cell surface (see Movie 4). **C.** Three CEV on the cell surface seen as a side view of the cell edge (see Movie 5). A different section of the same tomogram has previously been shown in Figure 7 and S4 (Hernandez-Gonzalez et al., 2023). **D.** Mid-view quantification of the percentage of CEV envelope in contact with the plasma membrane (left) and the distance between the CEV envelope and the plasma membrane (right). **E.** Left image correspond to the magenta boxed region in C, while the yellow arrow indicates a 13.75-nm wide linescan from which the greyscale values were plotted to identify the densities corresponding to the indicated structures from the virus core interior to cytoplasm (bottom left). The right images show labelling of these structures.

In addition to intact CEV, there were also many extracellular vesicles and membrane fragments as well as a surprising proportion of virions with broken membranes on the outside of the plasma membrane (35.2% CEV/EEV (extracellular enveloped virus) with fragmented outer envelope, n = 91; Figure 3 and Movie 6). It was not always obvious in the tomograms whether virions were still associated with the cell surface or were EEV that had been released from the cell, including multiple virions within a single envelope. To further confirm CEV frequently have a disrupted outer membrane, we performed immunofluorescence analysis in the absence of membrane permeabilization using antibodies against A27, an IMV surface protein, which would be inaccessible in intact CEV. In agreement with our tomograms and previous studies (Husain et al., 2007; Yu et al., 2026), we found that a considerable proportion of B5 positive CEV were also labelled with anti-A27 antibodies (Figure 3D).

**Figure 3.**
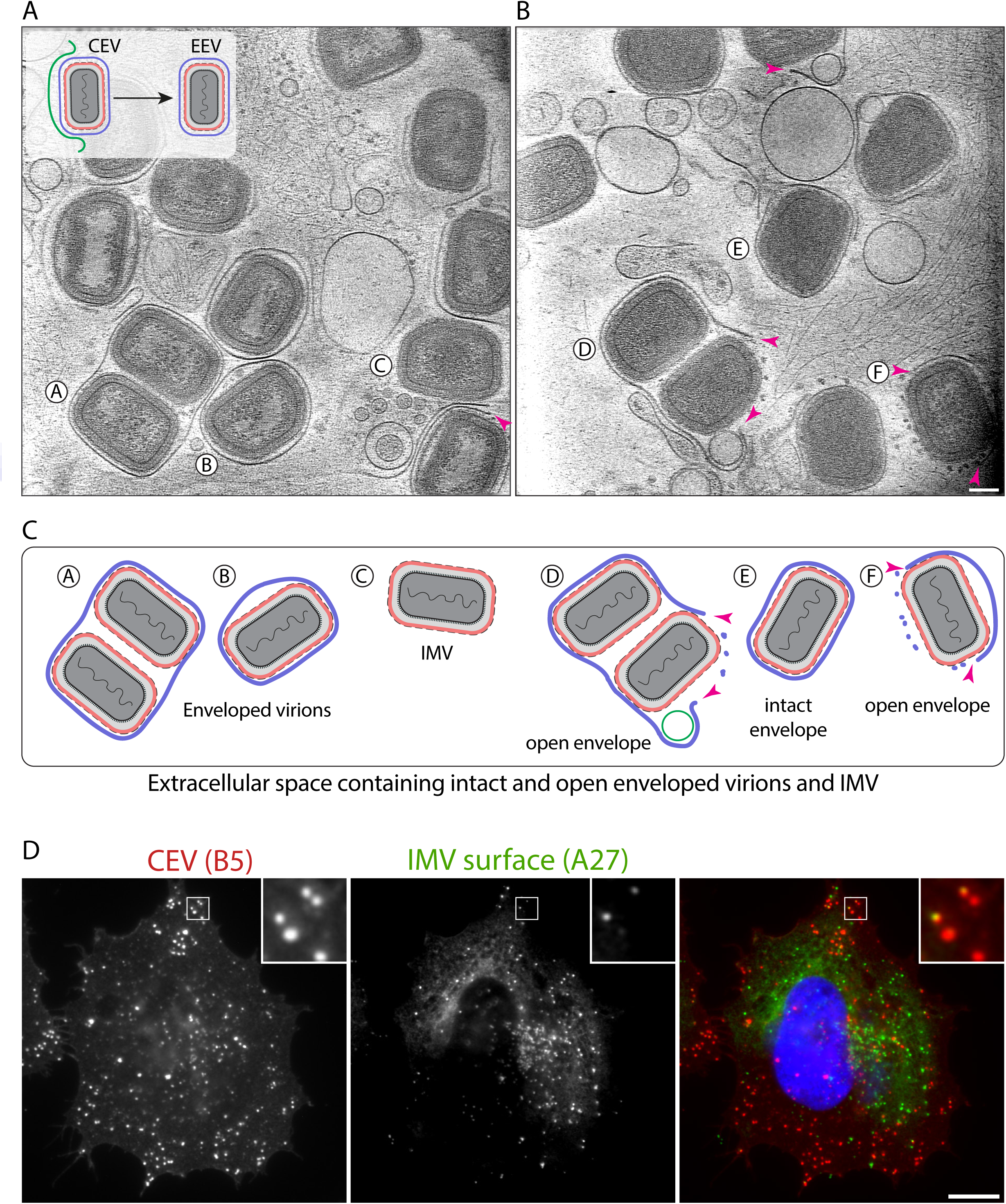
Extracellular virions and their broken envelopes. **A** and **B.** Tomogram sections showing extracellular viruses and their envelopes (see Movie 6). We quantified that 35.2% of CEV/EEV have a fragmented outer envelope as determine by tomogram examination of 91 viral particles in situ. **C.** Schematic representation of the virions labelled with letters in A and B. In all cases arrowheads point at the ends of broken membranes. **D.** Representative immunofluorescence image of non-permeabilized infected cell showing extracellular CEV positive for B5 and the IMV surface protein A27 at 8 hours post infection. Scalebar = 10 µm.

### Septin-like filaments beneath CEV

In the absence of virus induced actin polymerization, the proportion of CEV recruiting septins increases dramatically from ∼ 14 to 85 % (Figure 4A) (Pfanzelter et al., 2018). Therefore, using a virus strain that does not induce actin polymerization increases the chance of detecting septin filaments beneath CEV. This is particularly important as septins have been challenging to visualise in yeast using cryo-ET (Bertin et al., 2012). Our previous analysis suggested clathrin fully coats the membrane beneath CEV (Humphries et al., 2012). To further enhance our ability to identify septins, we collected tomograms on cells infected with the A36-YdF ΔNPF virus, which, as well as not assembling actin, is also defective in clathrin recruitment (Snetkov et al., 2016). We examined the surroundings of the plasma membrane beneath CEV to identify any structures that could represent septin filaments. In several tomograms that were of high quality, we found filamentous structures beneath CEV (Figures 4 and S3). These filaments are more easily observed in regions that lack actin filaments, such as blebs (Figure 4B and Movie 7). The filaments were ∼4-nm wide, which is consistent with the structure of single septin filaments visualized in vitro using cryo-EM (Beber et al., 2019; Lobato-Marquez et al., 2021; Mendonca et al., 2021). These filaments, whose ends often appeared frayed, extend across the span of the curvature formed by the virus at the plasma membrane albeit at different planes in the tomogram (Figures 4C and S3A and Movie 8). The putative septin filaments were recruited to CEV-plasma membrane domains of different curvatures and dimensions (Figures 4C and S3A and Movie 8). These filaments do not directly contact the plasma membrane and always run parallel to the membrane at a uniform distance of 21.15 ± 0.71 nm (Figures 4D and 5A-B). In addition, these putative septin filaments are noticeably different from actin or intermediate filaments, when these are compared from the same tomograms (Figure 5C). Finally, we could see possible connections between the putative septin filaments and the plasma membrane (Figure 4B). We consider it unlikely that these connections are actin, since septins are still recruited beneath CEV in Latrunculin B-treated cells (Figure S4).

**Figure 4.**
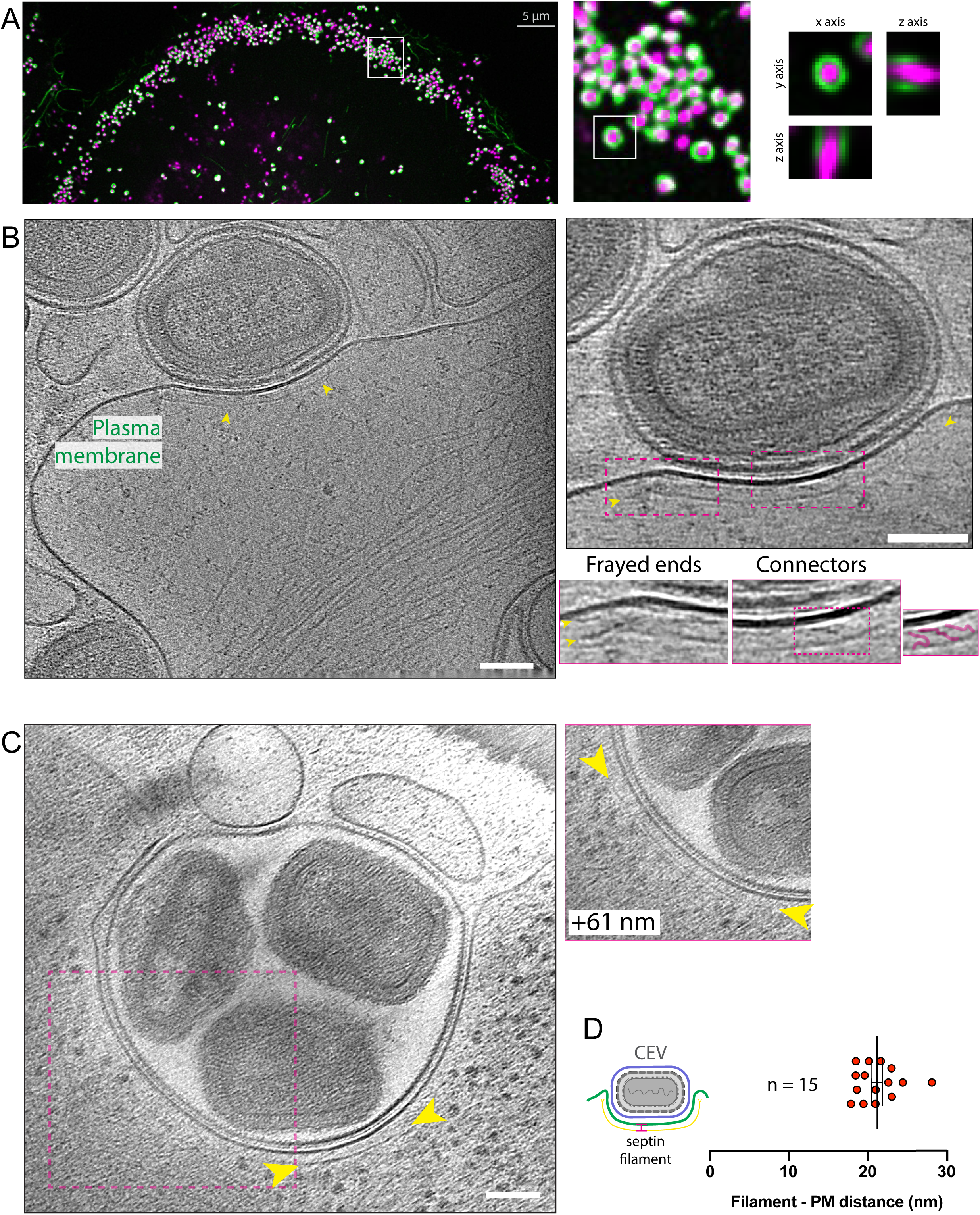
Septin filaments recruited to CEV. **A.** Maximum intensity project images still showing the recruitment of GFP-Septin 6 to CEV in live cells infected with the RFP-A3 A36-YdF vaccinia virus for 8 hours. In the images RFP-A3 viral core protein (magenta) and GFP-Septin 6 (green). Scalebar in main panel = 5 µm. **B.** The left image is a cryo-tomogram mid-section showing the presence of a thin filament to the curved plasma membrane beneath a CEV. The right panels show close-ups highlighting the frayed ends of the filament (yellow arrow heads) and putative connectors (magenta) between the septin filament and the plasma membrane. (See Movie 7). **C.** Tomogram section shows the recruitment of a septin filament (yellow arrow heads) beneath a CEV containing three virions on the cell surface. The filament extends around the curvature of the CEV in different planes of the tomogram. (See Movie 8). **D.** The graph shows the quantification of the distance between of septin filament and plasma membrane from 15 CEV tomograms. All cryo-ET scalebars = 100 nm.

**Figure 5.**
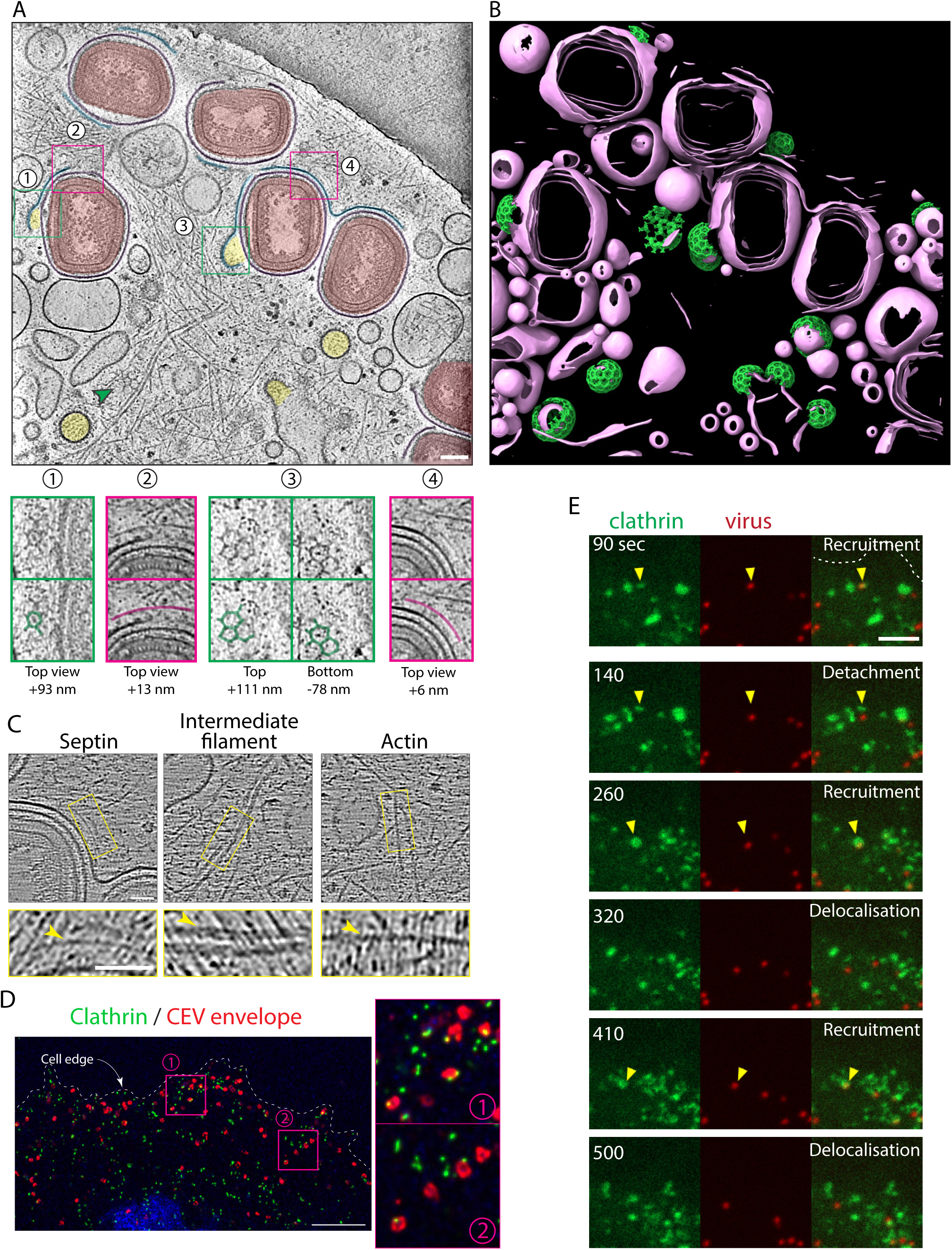
Structure and dynamics of clathrin recruited to CEV. **A.** Top-down tomogram view of CEV on a cell showing clathrin-associated vesicles and membranes (lumen in yellow), CEV (red and purple envelope) and plasma membrane ridges (blue). The images below correspond to the numbered squares in the main panel with or without clathrin (green) or septin (magenta) labelling. Scalebar = 100 nm. (See Movie 9). **B.** Automatic membrane segmentation using MemBrain (pink) and clathrin-AP2 template matching (green) using Pytom. **C.** Putative septin, intermediate and actin filaments from tomogram in A. Scale bar = 50 nm. **D.** Maximum intensity structured illumination microscopy image showing clathrin (green) localises in subdomains beneath B5 labelled CEV (red). Scalebar = 10 µm. **E.** Structured illumination microscopy image stills at the indicated times of a live cell infected with the RFP-A3 A36-YdF virus reveal multiple rounds of clathrin (green) recruitment to a CEV (red). (See Movie 10). Scalebar = 1 µm.

### Clathrin organisation beneath CEV

Consistent with its role in endocytosis, we could visualise clathrin honeycomb lattices decorating the plasma membrane and intracellular vesicles in tomograms of A36-YdF vaccinia infected cells (Figure 5A and Movie 9). In addition, we observed clathrin structures on the plasma membrane beneath CEV. We are confident that these structures are clathrin as they could be automatically identified by template matching using a single clathrin triskelion together with its adaptor AP-2 bound to the plasma membrane (Figure 5B). Furthermore, when copies of the given structure are back plotted into the specific positions at the tomogram, they are arranged with the spacing and polyhedral organisation of clathrin coats. Clathrin did not fully cover the plasma membrane beneath the CEV, but rather formed clathrin coated invaginations in subdomains beneath the virion (Figure 5A and S5). These clathrin subdomains have similar dimensions to those formed during clathrin-mediated endocytosis. When CEV also recruit putative septin filaments, these were located outside the clathrin subdomains (Figure 5A and S3A). Our cryo-EM observations of septin and clathrin are consistent with the co-recruitment of clathrin adaptor AP-2 and septin to ∼5% of CEV we previously determined using immunofluorescence microscopy (Pfanzelter et al., 2018). In addition, clathrin can deform not only the plasma membrane but also the attached CEV envelope, suggesting strong connections exist between the two membranes (Figure S6). Our previous immunofluorescence analysis did not show clathrin subdomains presumably because of the limited resolution of spinning disk confocal microscopy (Humphries et al., 2012). However, using instant structured-illumination microscopy (iSIM) we were able to observe clathrin subdomains on CEV in A36-YdF virus infected cells, consistent with our cryo-ET data (Figure 5D). The clathrin pits beneath CEV exhibit varying degrees of invagination (Figures 5A and S6), consistent with clathrin mediated endocytosis-like progression. Live cell iSIM imaging also revealed clathrin is recruited multiple times to the same virus suggesting that multiple endocytic events can occur beneath CEV on the plasma membrane (Figure 5E and Movie 10).

## Discussion

While we know the main viral proteins required, we still lack a structural understanding of the processes involved in generating the different virion types that the poxvirus infection produces. This includes the envelopment of IMV by a modified Golgi/endosomal cisterna to form IEV, which is required for exocytosis of the virus from the cell in the absence of lysis, a process critical for the spread of infection. Previous electron microscopy of thin sections of fixed infected cells reveals two-dimensional images of membrane cisternae enveloping IMV (Blasco and Moss, 1991; Cepeda and Esteban, 2014; Schmelz et al., 1994; Sodeik et al., 1993; Tooze et al., 1993). Images in thin sections, however, do not reveal the full complexity of IMV wrapping in three dimensions. By performing cryo-ET on vaccinia infected cells, we have now obtained three-dimensional insights into the process of IEV formation that are also free from fixation, processing and staining artifacts.

Collectively, our tomograms are consistent with a progressive zippering-like process mediated by the close association of the wrapping membrane with the IMV surface. We only observe single membrane cisterna associating with IMV, some of which are too small to fully wrap the virus. Hypothetically, these might be either dead-end events, or they might achieve full IMV envelopment by the binding and fusion of additional membrane cisternae or vesicles, or by cisternal growth via lipid transfer events. At the other extreme, there are cases where there is excess membrane during envelopment. Even when excess wrapping membrane is present, the inner membrane is still tightly associated with the IMV surface. A “loose” outer membrane might allow the membrane cisterna to more easily flow around the virus as its inner surface associates with the IMV in a zippering-like process. The fact that we observe CEV containing twin and triplet IMV suggests the membrane can envelope multiple virions *en masse*. The “loose” outer membrane also suggests that the interaction of viral proteins across the lumen of the membrane cisterna are not required for wrapping per se. In addition, we observed constrictions in the outer membrane when excess wrapping cisternae was present. Moreover, collars of protein density were visible in some of these constrictions suggesting that membrane remodelling is occurring. The removal of excess membrane during the wrapping process would explain why IEV in general do not have large amounts of excess outer membrane. Our observations, suggest that IMV envelopment is a flexible and adaptable process, which has many parallels to phagophore formation and engulfment of intracellular material during autophagy (Bieber et al., 2022; Dikic and Elazar, 2018). This flexibility may also explain why the enveloping membrane can tolerate variable amounts of transmembrane viral proteins such as A36 (Basant and Way, 2023).

Another striking feature we occasionally observed is the presence of a pore or aperture in the enveloping cisterna that are fully wrapped around the IMV. Because a fully sealed IEV is a topological requirement for the virus to undergo exocytosis at the plasma membrane, this pore must be sealed, for the wrapping cisternae to physically separate into the outer and inner IEV membranes, the latter of which is in contact with the IMV surface. We believe sealing the pore represents the last stage of IEV formation. Given the topology, we hypothesise that this pore is sealed by ESCRT-dependent annular fusion (Gatta and Carlton, 2019; Hernandez-Gonzalez et al., 2021; Remec Pavlin and Hurley, 2020). Consistent with an involvement of ESCRT, previous observations have demonstrated that the knockdown of TSG101 (ESCRT I complex) and the ESCRT-associated protein Alix results in a reduction of extracellular enveloped virions (Honeychurch et al., 2007; Huttunen et al., 2021). In addition, the vaccinia protein F13, which localises on the enveloping cisternae and is essential for IEV formation (Blasco and Moss, 1991) contains a putative ESCRT-recruiting late domain (YxxL) which could interact with Alix and is required for the formation of extracellular enveloped virions (Honeychurch et al., 2007). Finally, ESCRT-III and VPS4B have been recently shown to play a role in extracellular enveloped virion formation and proposed to mediate IMV envelopment by multivesicular bodies (Huttunen et al., 2021). Based on our observations, however, we favour a model by which all IMV envelopment events involve ESCRT-mediated sealing. The detection of ESCRT components on membrane cisterna enveloping IMV may be challenging given the transient nature of the sealing process, which is likely to be rapid.

We found that, after IEV fuse with the plasma membrane, CEV are closely associated with the plasma membrane beneath them, which was the outer IEV membrane prior to exocytosis (Figure 1C). This is further supported by Figure S6, showing clathrin can distort both the plasma membrane and CEV membrane. This close association must be mediated by interactions between viral proteins on the outside surface of the plasma membrane and the CEV. This contrasts the situation inside the cell where the outer and inner membranes of IEV as well as the cisterna enveloping IMV are not always closely opposed. This difference may in part reflect the oxidising environment of the extracellular space. In agreement with previous work, we also observed that the CEV envelope not in contact with the plasma membrane is often broken, which has important implications for immune recognition and virus entry (Husain et al., 2007).

To overcome the transient nature of septin recruitment to CEV, we used an A36-YdF mutant strain that cannot polymerise actin, leading to ∼80% of CEV recruiting septins (Pfanzelter et al., 2018). The use of this strain was essential for our work and shows how combining mutant conditions with cryo-ET can shed light on critical albeit transient structural arrangements. Using this virus, we observed ∼4 nm thin filaments that extend across the plasma membrane beneath CEV by cryo-ET, which is consistent with the width of septin filaments imaged in vitro by cryo-electron microscopy (Beber et al., 2019; Lobato-Marquez et al., 2021; Mendonca et al., 2021). In addition, their appearance is different from intermediate or actin filaments (Figure 5C). The filaments are present on different curvatures, which is in line with the broad range of curvatures septins can bind to (Cannon et al., 2017; Curtis and Gladfelter, 2024). Moreover, filaments vary in length, which might be determined by the contact area between the CEV and the plasma membrane. In addition, the observed frayed ends suggest septin filament ends might be organised differently from the rest of the filament.

Curiously, the putative septin filaments we see are at a constant distance (21.15 ± 0.71 nm) from the plasma membrane. In some cases, we can also see connecting densities between the filament and the plasma membrane, which do not appear to form straight, orthogonal links between filament and membrane (Figure 4B). It might be possible one of the IEV proteins exposed on the inner leaflet of the plasma membrane correspond to these connectors. However, the IEV proteins are too small to be these connections (Smith et al., 2002; Vernuccio et al., 2025) or big enough but not required to recruit septins, as in the case of A36 (Figure 4.7 in (Pfanzelter, 2018)). These connections are also unlikely to be actin, as septins are still present beneath CEV in Latrunculin B-treated cells (Figure S4). Strikingly, septin filaments are also a constant distance of ∼20 nm from the membrane at the bud neck in yeast (Bertin et al., 2012). In addition, cryo-ET of recombinant septin hexamers bound to *Mycobacterium smegmatis* and *Shigella flexneri* in vitro reveals they are spaced ∼16 and ∼20 nm from the bacterial surface, respectively (Lobato-Marquez et al., 2021). This shared feature further supports the identity of our filaments and suggests that the constant spacing is mediated by the septins themselves (i.e. the connectors are septins). This immediately raises a question: what are septins binding to on the plasma membrane beneath CEV? The most likely candidate is the membrane itself given the known properties of septins (Bridges et al., 2014; Cannon et al., 2017). However, this does not explain why septins are not recruited to IEV whose surface is, in principle, the same as the plasma membrane beneath CEV.

It is difficult to conceive how a single septin filament would be able to suppress the release of CEV from the cell surface (Pfanzelter et al., 2018). In structured illumination microscopy images, septins appear to form a ring around the majority of CEV in the absence of actin polymerization (Figure 4A) (Pfanzelter et al., 2018). However, in our tomograms in side views, we observe putative septin filaments at the plasma membrane directly beneath CEV (Figures 4 and S3). This difference suggests that septins beneath CEV are not detected by SIM because of the limited resolution in the Z axis. In support of this notion, B5, an integral membrane protein in the CEV envelope and in the plasma membrane beneath it, also appears as a ring in SIM imaging (Figure 3.11 in (Pfanzelter, 2018)). The lack of B5 signal at the top or bottom of the CEV confirms the limitations of SIM imaging and partly explains the differences seen between SIM and cryo-ET. Consistent with SIM imaging, we have also found putative septin filaments around CEV in top-down views by cryo-ET (Figure 5A). In both side and top-down views, we only visualise single septin filaments positioned perpendicular to the electron beam, at which the loss of information due to the missing wedge is minimal (Figures 4 and S3). This would suggest that we are only seeing a subset of the septin filaments that are recruited to CEV. Considering the information obtained from SIM and cryo-ET imaging modalities we favour a model by which multiple septin filaments are recruited to CEV, where they form a cage around the plasma membrane where the CEV resides.

We previously demonstrated that three NPF motifs in A36 promote the recruitment of AP2 (Snetkov et al., 2016), the adaptor responsible for recruiting clathrin to the plasma membrane, which promotes clustering of A36 beneath CEV prior to viral induced actin polymerisation (Humphries et al., 2012). This clustering helps polarise N-WASP beneath CEV, enhancing its stability and leading to more efficient and sustained Arp2/3-mediated actin polymerization. Consequently, in the absence of AP-2 recruitment, actin tail formation takes around twice as long (Humphries et al., 2012). However, the role of clathrin in the organisation and polarisation of A36 remained unclear. Our current study reveals, via cryo-ET and SIM, that clathrin forms membrane invaginations at subdomains beneath CEV that are indistinguishable from canonical clathrin-coated pits. These subdomains cover a small part of the plasma membrane beneath the CEV rather than the whole membrane beneath the virus. This finding might help explain the mechanism by which clathrin enhances actin polymerisation beneath the CEV. In yeast, Arp2/3 driven actin polymerisation is required to generate force to overcome the turgor pressure across the plasma membrane during clathrin-mediated endocytosis (CME) (Basu et al., 2014 ; Lu et al., 2016). In contrast, in mammalian cells, actin is only required for CME under conditions of increased membrane tension (Aghamohammadzadeh and Ayscough, 2009; Boulant et al., 2011; Gottlieb et al., 1993; Kaksonen et al., 2006; Serwas et al., 2022). The tight association of the CEV and plasma membrane over a large surface area suggests that Arp2/3-mediated actin polymerisation is similarly required for CME beneath CEV. We propose that actin polymerisation induced by CME at the relatively small subdomain beneath the CEV provides a template for subsequent viral-induced actin assembly downstream of A36 phosphorylation and N-WASP recruitment. Additionally, clathrin may locally concentrate phosphorylated A36 at the subdomains it forms, which would help amplify the signalling cascade that drives sustained Arp2/3-dependent actin polymerization. Altogether, we propose a model in which the actin polymerisation occurring during CME functions to enhance the ability of CEV to induce actin polymerization required to propel the virion away from the cell, thereby promoting the spread of infection. At the same time, endocytic recycling of viral proteins via CME back to the Golgi helps sustain IMV envelopment and A36 dependent kinesin-based transport of IEV throughout the many hours of infection (Husain and Moss, 2003; Husain and Moss, 2005; Ward and Moss, 2000). The acquisition of endocytic NPF motifs by A36 during evolution, and therefore its ability to recruit clathrin, likely conferred a dual selective advantage, by promoting its recycling and by enhancing actin polymerisation beneath CEV.

We have characterised vaccinia egress using a combination of cryo-ET and structured illumination microscopy of infected cells. We provide new insights into envelopment, and IEV/CEV architecture as well as showing the organisation of clathrin and putative septin filaments beneath CEV. Future work will be necessary to definitely confirm the filaments we observe are septins and to determine how they are recruited to CEV. Likewise, the detection of ESCRT components on enveloping IMV during pore sealing is an interesting future avenue to explore.

## Methods

### Cell growth, vaccinia infection and vitrification

The cryo-ET analysis reported in this article draws on multiple datasets, some of which were also analysed in a previous article (Hernandez-Gonzalez et al., 2023). Briefly, HeLa cells grown in complete MEM (with 10% fetal bovine serum, 100 ug/ml streptomycin, and 100 U/ml penicillin) at 37 °C with 5% CO_2_, were washed with PBS, incubated with trypsin, seeded on glow-discharged Quantifoil R3.5/1 or R2/2 gold grids of 200 mesh (40 s at 45 mA) and incubated in 6-well plates with complete MEM overnight. The cells were subsequently infected with sucrose-purified A36-YdF ΔF11, A36-YdF ΔNPF1-3 ΔF11 or ΔF11 vaccinia strains in serum free MEM at a multiplicity of infection of 2 (Albarnaz et al., 2023; Cordeiro et al., 2009; Hernandez-Gonzalez et al., 2023). One hour post-infection (hpi), the medium was replaced with complete MEM. At 8 hpi, grids were washed with PBS and Whatman paper was used to remove excess PBS before automatic blotting and vitrification using Vitrobot Mark IV System, at 95% relative humidity and 20°C. 10-nm colloidal gold particles were pipetted onto grids ahead of blotting for 14s with a relative force of -10.

### Cryo-electron tomography, image processing and analysis

A Talos Arctica or Glacios TEM (Thermo Fisher) were used to screen grids and select those with optimal confluency and cellular thinness. Grid regions were recorded before transferring to a Titan Krios TEM (Thermo Fisher), where tilt series were acquired. We acquired 187, 238 and 126 tilt series for cells infected with ΔF11, A36-YdF ΔF11 and A36-YdF ΔNPF1-3 ΔF11, respectively. The Titan Krios was fitted with either a K2 Summit camera with a Bioquantum energy filter (Gatan) or a Falcon 4i camera with a Selectris energy filter (Thermo Fisher). Dose-symmetric tilt series were collected from −57° to +57° at a 3° increment, a pixel size of 4.31 or 3.74 Å, defocus of −8 μm or -7 μm and dose 1.7 or 2.01 e/ Å^2^ per tilt, respectively, using Tomography software (Thermo Fisher). Four movie frames were collected per tilt. Alignframes from IMOD was used to align movie frames (Kremer et al., 1996). Tilt series alignments using patchtracking or gold fiducials, contrast transfer function (CTF) correction, and tomogram reconstruction were all conducted in IMOD (Kremer et al., 1996; Xiong et al., 2009). Typically, tomograms were reconstructed using 3× binning, yielding voxel sizes of 11.22 or 1.93 Å for tilt series acquired at pixel sizes of 4.31 or 3.74 Å, respectively. Manual segmentations were performed using AMIRA (Thermo Fisher) after denoising using ISONet with CTF deconvolution and missing wedge filling from deep learning (Liu et al., 2022). The distance between septin filaments and plasma membrane as well as between the CEV outer membrane and plasma membrane were measured using 3dmod from IMOD at the mid-plane of each CEV (Kremer et al., 1996). Three measurements per event were averaged to obtain the final value. To determine the percentage of CEV outer membrane in contact with the plasma membrane, contours of both membranes were traced in 3dmod, and their lengths used to calculate such percentage.

### Clathrin template matching and automatic membrane segmentation

We ran template matching using Pytom (Chaillet et al., 2023) to identify clathrin vertices in tomograms, using an EM map of a clathrin–AP-2 complex on the plasma membrane of human HSC-3 cells (EMDB-46973) (Sun et al., 2025). By then placing the template at the discovered positions and orientations in the tomogram, they together assembled a clathrin coat consistent with previous structure of clathrin cages, for example in the size of clathrin hexagons and pentagons. Automatic membrane segmentation shown in Figure 5B was performed using MemBrain with default parameters (Lamm et al., 2022).

### Immunofluorescence analysis and live cell imaging

HeLa cells grown on fibronectin-coated coverslips were infected with the A36-YdF virus (Rietdorf et al., 2001) for 8 hours and fixed with 4% paraformaldehyde in PBS for 15 min, incubated in blocking buffer (10 mM MES (pH 6.1), 150 mM NaCl, 5 mM EGTA, 5 mM MgCl_2_, and 5 mM glucose) containing 2% (v/v) fetal calf serum and 1% (w/v) BSA for 30 min before incubation with a rat monoclonal antibody against B5 (19C2) (Schmelz et al., 1994) for an hour, followed by incubation with a goat Alexa Fluor 647 anti-rat secondary antibody (Invitrogen; 1:1000). To determine antibody accessibility to the IMV surface, a C3 monoclonal antibody against A27 was used (1:1000; (Rodriguez et al., 1985)) followed by a donkey Alexa Fluor 488 anti-mouse secondary antibody (1:1000, Invitrogen) for 1 h each. Finally, cells were permeabilised with 0.1% Triton-X100/PBS for 5 min and stained with 4,6-diamidino-2-phenylindole (DAPI) for 5 min, before mounting using Mowiol (Sigma). Coverslips were imaged on a Zeiss Axioplan2 microscope with a 63x/1.4 NA Plan-Achromat objective and a Photometrics Cool Snap HQ camera, controlled by MetaMorph. The experiment was repeated 3 times and, in each case, at least 10 cells were imaged. Fiji was used for image analysis.

To study the effect of Latrunculin B on septin recruitment to CEV, A36-YdF virus infected HeLa cells on fibronectin-coated coverslips were treated with DMSO control or 5 µM Latrunculin B (Sigma) for 5 min before fixation. Coverslips were then incubated with blocking buffer, anti-B5 antibody, and anti-Rat 647-alexa conjugated antibody as above, before permeabilization with 0.1% Triton-X100/PBS 5 min. Septin 7 was labelled with a rabbit polyclonal antibody against septin7 (1:300, 18991 IBL) followed by an Alexa Fluor 488 anti-rabbit secondary antibody (1:1000, Thermo Fisher) for 1 h prior to DAPI staining and Mowiol mounting. Coverslips were imaged by Structured Illumination Microscopy (VT-iSIM). To image septin and vaccinia in live cells, a GFP-Septin 6 HeLa cell line (Pfanzelter et al., 2018) was grown on fibronectin-coated Mattek dishes overnight, before infection with an RFP-A3 A36-YdF vaccinia strain (Xu et al., 2023) and imaged 8 hpi under VT-iSIM. VT-iSIM was conducted on an Olympus iX83 Microscope with Olympus 150x/1.45 NA X-Line apochromatic objective lens, dual Photometrics BSI-Express sCMOS cameras, and CoolLED pE-300 light source (Visitech), controlled using Micro-Manager. Image z-stacks with 100 nm steps were acquired and deconvolved using express deconvolution on Huygens Software (Scientific Volume Imaging). Point spread function measurement of sub-diffraction 100-nm beads confirmed an XY resolution of 125 nm. To visualise clathrin dynamics beneath CEV, HeLa cells were infected with the RFP-A3 A36-YdF vaccinia virus and transfected 2 hours later with a 1 μg of a plasmid expressing GFP-LCa (clathrin light chain) under the control of viral promoter pEL (Humphries et al., 2012) using FUGENE as described by the manufacturer (Promega). At 8 hpi, Mattek dishes were transferred to the VT-iSIM microscope at 37°C, and images were acquiring at 10-second intervals for 50 time points (total duration: 8 min 10 s) in a single z-plane.

To visualise clathrin localisation to CEV, HeLa cells were infected with a A36-YdF virus strain, fixed 8 hpi, incubated with blocking buffer prior to TritonX100 0.1% / PBS. and stained with rabbit polyclonal antibody against clathrin heavy chain (1:500, ab21679 Abcam) and donkey Alexa Fluor 555 anti-rabbit secondary antibody (1:1000, Thermo Fisher). Finally, coverslips were stained with DAPI and mounted using Mowiol and images acquired on the VT-iSIM microscope.

## Supporting information

All supplemental figures

## Acknowledgements

We thank Angika Basant and Jeremy Carlton (King’s College London) as well as Serge Mostowy (The London School of Hygiene and Tropical Medicine) for insightful comments on the manuscript. We also acknowledge Philip Walker and Andy Purkiss of the Structural Biology Science Technology Platform and the Scientific Computing Science Technology Platform for computational support.

## Author contributions

All authors designed the research program and strategy. MHG, TC and AN performed research, MHG and TC analysed data, MHG and MW wrote the paper and all authors provided feedback on the manuscript.

## Supplementary figure legends

**Figure S1. Cryo-ET gallery of vaccinia envelopment.**

**A.** Tomograms showing examples of IMV envelopment. Labelled images indicate the enveloping membranes in blue and arrowheads point at dilated cisternae ends. **B.** Different tomogram sections of the same envelopment showing 3 different pores marked with numbers and magenta arrowheads (see Movie 2). **C.** Envelopment next to a fully sealed IEV. In both cases the envelope is in close contact to microtubules (red). **D.** Example of envelopment around an aberrant IMV. We have characterised a total of 12 wrapping events from our tomograms.

**Figure S2. Cryo-ET gallery of IEV.**

**A.** Tomogram sections showing middle views of IEV. The magenta arrowheads indicate an example where the inner IEV membrane is broken. **B.** Two different sections of a tomogram showing an IEV with part of its inner membrane broken (magenta arrowheads) and possible remodelling of the out IEV membrane (green arrowhead). **C.** CEV containing three IMV. Scalebar = 100 nm.

**Figure S3. Cryo-ET gallery of septin filaments beneath CEV on the plasma membrane.**

**A.** Pairs of tomogram sections showing septin filament recruitment (magenta arrowheads) to CEV on the plasma membrane. Different parts of the same filament are seen in the two z-sections. The example at the bottom shows a clathrin coated invagination (orange arrowhead) recruited next to a septin filament. **B.** Middle section of CEV demonstrates septin filaments can be recruited to a variety of curvatures. Scalebar = 100 nm.

**Figure S4. Septin recruitment is independent of actin polymerization** Maximum intensity projection structured illumination microscopy images showing that inhibition of actin polymerization with Latrunculin B (LatB) does not impact the recruitment of Septin7 (green) to CEV (Red labelled with B5). Scalebar = 10 µm.

**Figure S5. Raw cryo-ET sections from Figure 5**. Tomogram sections at similar positions as Figure 5A and + 111 nm close-up, in the absence of ISONet denoising. Binned 3, SIRT-like filtered, and CTF corrected. Scalebar = 100 nm.

**Figure S6. Cryo-ET sections of clathrin deforming both the plasma membrane and CEV envelope.** Green arrowheads point at clathrin-induced invaginations, while magenta squares highlight the clathrin hexagonal lattice recruited to the invaginations. Scalebar = 100 nm.

## Movie legends

**Movie 1. IMV envelopment and final pore.**

Tomogram corresponding to the section shown in Figure 1A on the right. Arrowhead points at the cisternal pore. Scalebar = 100 nm.

**Movie 2. Envelopment case with multiple pores.**

Three Tomogram showing IMV envelopment and three pore at the enveloping cisterna. Section of this tomogram are shown in Figure S1B. Scalebar = 100 nm.

**Movie 3. Membrane remodelling during IMV envelopment.**

Tomogram showing remodelling (tubulation and constriction) of the enveloping cisterna. Red arrow points at a pore, while the green arrow points a membrane constriction displaying a structural collar around it. It corresponds to the image shown in Figure 1B. Scalebar = 100 nm.

**Movie 4. Top-down view of CEV on the cell surface.**

Tomogram of two CEV on the cell surface, as shown in Figure 2B. Scalebar = 100 nm.

**Movie 5. Side view of CEV on the cell surface.**

Three CEV on the cell surface corresponding to the side view shown in Figure 2C.

**Movie 6. Extracellular virions and their broken envelopes.**

Tomogram from the section shown in Figure 3A. The planes that mostly show the cell interiors are labelled (white), while the planes showing extracellular virions have a yellow label.

**Movie 7. Septin filament beneath CEV.**

Tomogram of the section shown in Figure 4B. Scalebar = 100 nm.

**Movie 8. Septin filament beneath CEV containing three IMV.**

Tomogram of the CEV particle shown in Figure 4C. Scalebar = 100 nm.

**Movie 9. Segmentation of CEV on the cell surface.**

Manual segmentation of tomogram corresponding to the image shown in Figure 5A.

**Movie 10. Multiple rounds of clathrin recruitment to CEV.**

Timelapse Movie corresponding to the stills showing in Figure 5E. Green channel is GFP-tagged clathrin light chain (GFP-LCa), while red signal corresponds to RFP-A3 viral core protein. Time is indicated in minutes and seconds.

## References

Aghamohammadzadeh, S., and K.R. Ayscough. 2009. Differential requirements for actin during yeast and mammalian endocytosis. Nat Cell Biol. 11:1039–1042.

Albarnaz, J.D., J. Kite, M. Oliveira, H. Li, Y. Di, M.H. Christensen, J.A. Paulo, R. Antrobus, S.P. Gygi, F.I. Schmidt, E.L. Huttlin, G.L. Smith, and M.P. Weekes. 2023. Quantitative proteomics defines mechanisms of antiviral defence and cell death during modified vaccinia Ankara infection. Nat Commun. 14:8134.

Basant, A., and M. Way. 2023. The amount of Nck rather than N-WASP correlates with the rate of actin-based motility of Vaccinia virus. Microbiol Spectr. 11:e0152923.

Basu, R., E.L. Munteanu, and F. Chang. 2014. Role of turgor pressure in endocytosis in fission yeast. Mol Biol Cell. 25:679–687.

Beber, A., C. Taveneau, M. Nania, F.C. Tsai, A. Di Cicco, P. Bassereau, D. Levy, J.T. Cabral, H. Isambert, S. Mangenot, and A. Bertin. 2019. Membrane reshaping by micrometric curvature sensitive septin filaments. Nat Commun. 10:420.

Beiras, C.G., E. Malembi, R. Escrig-Sarreta, S. Ahuka, P. Mbala, H.M. Mavoko, L. Subissi, A.B. Abecasis, M. Marks, and O. Mitja. 2025. Concurrent outbreaks of mpox in Africa-an update. Lancet. 405:86–96.

Bertin, A., M.A. McMurray, J. Pierson, L. Thai, K.L. McDonald, E.A. Zehr, G. Garcia, 3rd, P. Peters, J. Thorner, and E. Nogales. 2012. Three-dimensional ultrastructure of the septin filament network in Saccharomyces cerevisiae. Mol Biol Cell. 23:423–432.

Bieber, A., C. Capitanio, P.S. Erdmann, F. Fiedler, F. Beck, C.W. Lee, D. Li, G. Hummer, B.A. Schulman, W. Baumeister, and F. Wilfling. 2022. In situ structural analysis reveals membrane shape transitions during autophagosome formation. Proc Natl Acad Sci U S A. 119:e2209823119.

Blasco, R., and B. Moss. 1991. Extracellular vaccinia virus formation and cell-to-cell virus transmission are prevented by deletion of the gene encoding the 37,000-Dalton outer envelope protein. J Virol. 65:5910–5920.

Boulant, S., C. Kural, J.C. Zeeh, F. Ubelmann, and T. Kirchhausen. 2011. Actin dynamics counteract membrane tension during clathrin-mediated endocytosis. Nat Cell Biol. 13:1124–1131.

Bridges, A.A., H. Zhang, S.B. Mehta, P. Occhipinti, T. Tani, and A.S. Gladfelter. 2014. Septin assemblies form by diffusion-driven annealing on membranes. Proc Natl Acad Sci U S A. 111:2146–2151.

Cannon, K.S., B.L. Woods, and A.S. Gladfelter. 2017. The Unsolved Problem of How Cells Sense Micron-Scale Curvature. Trends Biochem Sci. 42:961–976.

Cavini, I.A., D.A. Leonardo, H.V.D. Rosa, D.K.S.V. Castro, H. D’Muniz Pereira, N.F. Valadares, A.P.U. Araujo, and R.C. Garratt. 2021. The Structural Biology of Septins and Their Filaments: An Update. Frontiers in Cell and Developmental Biology. 9.

Cepeda, V., and M. Esteban. 2014. Novel insights on the progression of intermediate viral forms in the morphogenesis of vaccinia virus. Virus Res. 183:23–29.

Chaillet, M.L., G. van der Schot, I. Gubins, S. Roet, R.C. Veltkamp, and F. Forster. 2023. Extensive Angular Sampling Enables the Sensitive Localization of Macromolecules in Electron Tomograms. Int J Mol Sci. 24.

Cordeiro, J.V., S. Guerra, Y. Arakawa, M.P. Dodding, M. Esteban, and M. Way. 2009. F11-mediated inhibition of RhoA signalling enhances the spread of vaccinia virus in vitro and in vivo in an intranasal mouse model of infection. PLoS One. 4:e8506.

Cudmore, S., P. Cossart, G. Griffiths, and M. Way. 1995. Actin-based motility of vaccinia virus. Nature. 378:636–638.

Curtis, B.N., and A.S. Gladfelter. 2024. Drivers of Morphogenesis: Curvature Sensor Self-Assembly at the Membrane. Cold Spring Harb Perspect Biol. 16.

Dikic, I., and Z. Elazar. 2018. Mechanism and medical implications of mammalian autophagy. Nat Rev Mol Cell Biol. 19:349–364.

Doceul, V., M. Hollinshead, L. van der Linden, and G.L. Smith. 2010. Repulsion of superinfecting virions: a mechanism for rapid virus spread. Science. 327:873–876.

Dodding, M.P., R. Mitter, A.C. Humphries, and M. Way. 2011. A kinesin-1 binding motif in vaccinia virus that is widespread throughout the human genome. EMBO J. 30:4523–4538.

Dodding, M.P., T.P. Newsome, L.M. Collinson, C. Edwards, and M. Way. 2009. An E2-F12 complex is required for intracellular enveloped virus morphogenesis during vaccinia infection. Cell Microbiol. 11:808–824.

Domi, A., A.S. Weisberg, and B. Moss. 2008. Vaccinia virus E2L null mutants exhibit a major reduction in extracellular virion formation and virus spread. J Virol. 82:4215–4226.

Engelstad, M., and G.L. Smith. 1993. The vaccinia virus 42-kDa envelope protein is required for the envelopment and egress of extracellular virus and for virus virulence. Virology. 194:627–637.

Frischknecht, F., V. Moreau, S. Rottger, S. Gonfloni, I. Reckmann, G. Superti-Furga, and M. Way. 1999. Actin-based motility of vaccinia virus mimics receptor tyrosine kinase signalling. Nature. 401:926–929.

Gatta, A.T., and J.G. Carlton. 2019. The ESCRT-machinery: closing holes and expanding roles. Curr Opin Cell Biol. 59:121–132.

Geada, M.M., I. Galindo, M.M. Lorenzo, B. Perdiguero, and R. Blasco. 2001. Movements of vaccinia virus intracellular enveloped virions with GFP tagged to the F13L envelope protein. J Gen Virol. 82:2747–2760.

Gottlieb, T.A., I.E. Ivanov, M. Adesnik, and D.D. Sabatini. 1993. Actin microfilaments play a critical role in endocytosis at the apical but not the basolateral surface of polarized epithelial cells. J Cell Biol. 120:695–710.

Gubser, C., S. Hue, P. Kellam, and G.L. Smith. 2004. Poxvirus genomes: a phylogenetic analysis. J Gen Virol. 85:105–117.

Hernandez-Gonzalez, M., T. Calcraft, A. Nans, P.B. Rosenthal, and M. Way. 2023. A succession of two viral lattices drives vaccinia virus assembly. PLoS Biol. 21:e3002005.

Hernandez-Gonzalez, M., G. Larocque, and M. Way. 2021. Viral use and subversion of membrane organization and trafficking. J Cell Sci. 134.

Hollinshead, M., G. Rodger, H. Van Eijl, M. Law, R. Hollinshead, D.J. Vaux, and G.L. Smith. 2001. Vaccinia virus utilizes microtubules for movement to the cell surface. J Cell Biol. 154:389–402.

Honeychurch, K.M., G. Yang, R. Jordan, and D.E. Hruby. 2007. The vaccinia virus F13L YPPL motif is required for efficient release of extracellular enveloped virus. J Virol. 81:7310–7315.

Humphries, A.C., M.P. Dodding, D.J. Barry, L.M. Collinson, C.H. Durkin, and M. Way. 2012. Clathrin potentiates vaccinia-induced actin polymerization to facilitate viral spread. Cell Host Microbe. 12:346–359.

Husain, M., and B. Moss. 2003. Intracellular trafficking of a palmitoylated membrane-associated protein component of enveloped vaccinia virus. J Virol. 77:9008–9019.

Husain, M., and B. Moss. 2005. Role of receptor-mediated endocytosis in the formation of vaccinia virus extracellular enveloped particles. J Virol. 79:4080–4089.

Husain, M., A.S. Weisberg, and B. Moss. 2007. Resistance of a vaccinia virus A34R deletion mutant to spontaneous rupture of the outer membrane of progeny virions on the surface of infected cells. Virology. 366:424–432.

Huttunen, M., J. Samolej, R.J. Evans, A. Yakimovich, I.J. White, J. Kriston-Vizi, J. Martin-Serrano, W.I. Sundquist, E.M. Frickel, and J. Mercer. 2021. Vaccinia virus hijacks ESCRT-mediated multivesicular body formation for virus egress. Life Sci Alliance. 4.

Kaksonen, M., C.P. Toret, and D.G. Drubin. 2006. Harnessing actin dynamics for clathrin-mediated endocytosis. Nat Rev Mol Cell Biol. 7:404–414.

Kremer, J.R., D.N. Mastronarde, and J.R. McIntosh. 1996. Computer visualization of three-dimensional image data using IMOD. J Struct Biol. 116:71–76.

Lamm, L., R.D. Righetto, W. Wietrzynski, M. Poge, A. Martinez-Sanchez, T. Peng, and B.D. Engel. 2022. MemBrain: A deep learning-aided pipeline for detection of membrane proteins in Cryo-electron tomograms. Comput Methods Programs Biomed. 224:106990.

Liu, Y.T., H. Zhang, H. Wang, C.L. Tao, G.Q. Bi, and Z.H. Zhou. 2022. Isotropic reconstruction for electron tomography with deep learning. Nat Commun. 13:6482.

Lobato-Marquez, D., J. Xu, G.O. Guler, A. Ojiakor, M. Pilhofer, and S. Mostowy. 2021. Mechanistic insight into bacterial entrapment by septin cage reconstitution. Nat Commun. 12:4511.

Lu, R., D.G. Drubin, and Y. Sun. 2016. Clathrin-mediated endocytosis in budding yeast at a glance. J Cell Sci. 129:1531–1536.

Makokha, G.N., M. Abuduwaili, K. Chayama, and M. Hijikata. 2025. What is the future of Mpox outbreak? Virology. 610:110618.

Mendonca, D.C., S.L. Guimaraes, H.D. Pereira, A.A. Pinto, M.A. de Farias, A.S. de Godoy, A.P.U. Araujo, M. van Heel, R.V. Portugal, and R.C. Garratt. 2021. An atomic model for the human septin hexamer by cryo-EM. J Mol Biol. 433:167096.

Moreau, V., F. Frischknecht, I. Reckmann, R. Vincentelli, G. Rabut, D. Stewart, and M. Way. 2000. A complex of N-WASP and WIP integrates signalling cascades that lead to actin polymerization. Nat Cell Biol. 2:441–448.

Moss, B. 2007. Poxviridae: the viruses and their replication. *In* Fields Virology. H.P. Knipe DM, editor. Lippincott Williams and Wilkins, Philadelphia. 2905–2946.

Mostowy, S., and P. Cossart. 2012. Septins: the fourth component of the cytoskeleton. Nat Rev Mol Cell Biol. 13:183–194.

Nakazawa, K., B. Chauvin, S. Mangenot, and A. Bertin. 2023. Reconstituted in vitro systems to reveal the roles and functions of septins. J Cell Sci. 136.

Newsome, T.P., N. Scaplehorn, and M. Way. 2004. SRC mediates a switch from microtubule- to actin-based motility of vaccinia virus. Science. 306:124–129.

Newsome, T.P., I. Weisswange, F. Frischknecht, and M. Way. 2006. Abl collaborates with Src family kinases to stimulate actin-based motility of vaccinia virus. Cell Microbiol. 8:233–241.

Pfanzelter, J. 2018. The role of septins during vaccinia virus spread. Vol. Ph.D. University College London.

Pfanzelter, J., S. Mostowy, and M. Way. 2018. Septins suppress the release of vaccinia virus from infected cells. J Cell Biol. 217:2911–2929.

Reeves, P.M., B. Bommarius, S. Lebeis, S. McNulty, J. Christensen, A. Swimm, A. Chahroudi, R. Chavan, M.B. Feinberg, D. Veach, W. Bornmann, M. Sherman, and D. Kalman. 2005. Disabling poxvirus pathogenesis by inhibition of Abl-family tyrosine kinases. Nat Med. 11:731–739.

Remec Pavlin, M., and J.H. Hurley. 2020. The ESCRTs - converging on mechanism. J Cell Sci. 133.

Rietdorf, J., A. Ploubidou, I. Reckmann, A. Holmstrom, F. Frischknecht, M. Zettl, T. Zimmermann, and M. Way. 2001. Kinesin-dependent movement on microtubules precedes actin-based motility of vaccinia virus. Nat Cell Biol. 3:992–1000.

Roberts, K.L., and G.L. Smith. 2008. Vaccinia virus morphogenesis and dissemination. Trends Microbiol. 16:472–479.

Rodriguez, J.F., R. Janeczko, and M. Esteban. 1985. Isolation and characterization of neutralizing monoclonal antibodies to vaccinia virus. J Virol. 56:482–488.

Rottger, S., F. Frischknecht, I. Reckmann, G.L. Smith, and M. Way. 1999. Interactions between vaccinia virus IEV membrane proteins and their roles in IEV assembly and actin tail formation. J Virol. 73:2863–2875.

Schampera, J.N., and C. Schwan. 2024. Septin dynamics and organization in mammalian cells. Curr Opin Cell Biol. 91:102442.

Schmelz, M., B. Sodeik, M. Ericsson, E.J. Wolffe, H. Shida, G. Hiller, and G. Griffiths. 1994. Assembly of vaccinia virus: the second wrapping cisterna is derived from the trans Golgi network. J Virol. 68:130–147.

Serwas, D., M. Akamatsu, A. Moayed, K. Vegesna, R. Vasan, J.M. Hill, J. Schoneberg, K.M. Davies, P. Rangamani, and D.G. Drubin. 2022. Mechanistic insights into actin force generation during vesicle formation from cryo-electron tomography. Dev Cell. 57:1132–1145 e1135.

Smith, G.L., A. Vanderplasschen, and M. Law. 2002. The formation and function of extracellular enveloped vaccinia virus. J Gen Virol. 83:2915–2931.

Snetkov, X., I. Weisswange, J. Pfanzelter, A.C. Humphries, and M. Way. 2016. NPF motifs in the vaccinia virus protein A36 recruit intersectin-1 to promote Cdc42:N-WASP-mediated viral release from infected cells. Nat Microbiol. 1:16141.

Sodeik, B., R.W. Doms, M. Ericsson, G. Hiller, C.E. Machamer, W. van’t Hof, G. van Meer, B. Moss, and G. Griffiths. 1993. Assembly of vaccinia virus: role of the intermediate compartment between the endoplasmic reticulum and the Golgi stacks. J Cell Biol. 121:521–541.

Spiliotis, E.T., and M.A. McMurray. 2020. Masters of asymmetry - lessons and perspectives from 50 years of septins. Mol Biol Cell. 31:2289–2297.

Sun, W.W., D.J. Michalak, K.A. Sochacki, P. Kunamaneni, M.A. Alfonzo-Mendez, A.M. Arnold, M.P. Strub, J.E. Hinshaw, and J.W. Taraska. 2025. Cryo-electron tomography pipeline for plasma membranes. Nat Commun. 16:855.

Tooze, J., M. Hollinshead, B. Reis, K. Radsak, and H. Kern. 1993. Progeny vaccinia and human cytomegalovirus particles utilize early endosomal cisternae for their envelopes. Eur J Cell Biol. 60:163–178.

Traub, L.M. 2011. Regarding the amazing choreography of clathrin coats. PLoS Biol. 9:e1001037.

van Eijl, H., M. Hollinshead, G. Rodger, W.H. Zhang, and G.L. Smith. 2002. The vaccinia virus F12L protein is associated with intracellular enveloped virus particles and is required for their egress to the cell surface. J Gen Virol. 83:195–207.

Vassilopoulos, S., and G. Montagnac. 2024. Clathrin assemblies at a glance. J Cell Sci. 137.

Vernuccio, R., A. Martinez Leon, C.S. Poojari, J. Buchrieser, C.N. Selverian, Y. Jaleta, A. Meola, F. Guivel-Benhassine, F. Porrot, A. Haouz, M. Chevreuil, B. Raynal, J. Mercer, E. Simon-Loriere, K. Chandran, O. Schwartz, J.S. Hub, and P. Guardado-Calvo. 2025. Structural insights into tecovirimat antiviral activity and poxvirus resistance. Nat Microbiol. 10:734–748.

Ward, B.M., and B. Moss. 2000. Golgi network targeting and plasma membrane internalization signals in vaccinia virus B5R envelope protein. J Virol. 74:3771–3780.

Ward, B.M., and B. Moss. 2001. Vaccinia virus intracellular movement is associated with microtubules and independent of actin tails. J Virol. 75:11651–11663.

Ward, B.M., and B. Moss. 2004. Vaccinia virus A36R membrane protein provides a direct link between intracellular enveloped virions and the microtubule motor kinesin. J Virol. 78:2486–2493.

Wolffe, E.J., S.N. Isaacs, and B. Moss. 1993. Deletion of the vaccinia virus B5R gene encoding a 42-kilodalton membrane glycoprotein inhibits extracellular virus envelope formation and dissemination. J Virol. 67:4732–4741.

Woods, B.L., and A.S. Gladfelter. 2021. The state of the septin cytoskeleton from assembly to function. Curr Opin Cell Biol. 68:105–112.

Xiong, Q., M.K. Morphew, C.L. Schwartz, A.H. Hoenger, and D.N. Mastronarde. 2009. CTF determination and correction for low dose tomographic tilt series. J Struct Biol. 168:378–387.

Xu, A., A. Basant, S. Schleich, T.P. Newsome, and M. Way. 2023. Kinesin-1 transports morphologically distinct intracellular virions during vaccinia infection. J Cell Sci. 136.

Yu, H., W. Resch, C.A. Cotter, W. Xiao, T. Karamanolis, A.A. Belghith, M.A. Ignacio, P.L. Earl, G.H. Cohen, and B. Moss. 2026. Immunogenicity of poxvirus A16/G9 entry-fusion subcomplex and its restriction by A56/K2 protein informs vaccine design. Nat Microbiol. 11:2008–2020.

