## Supplementary figures and images for "In situ cryo-electron tomography of vaccinia virus exit from infected cells"

### All supplemental figures

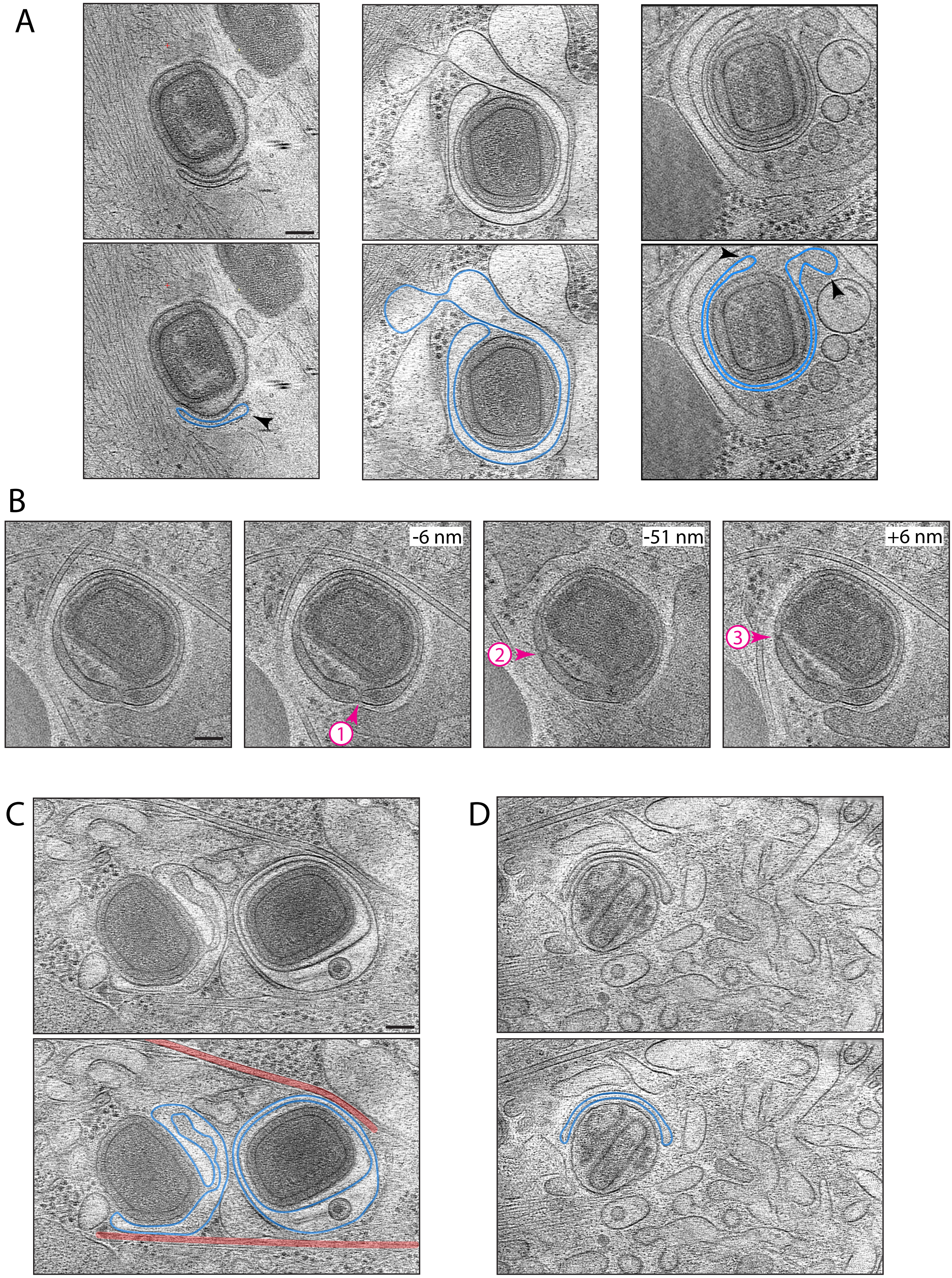


Figure S1

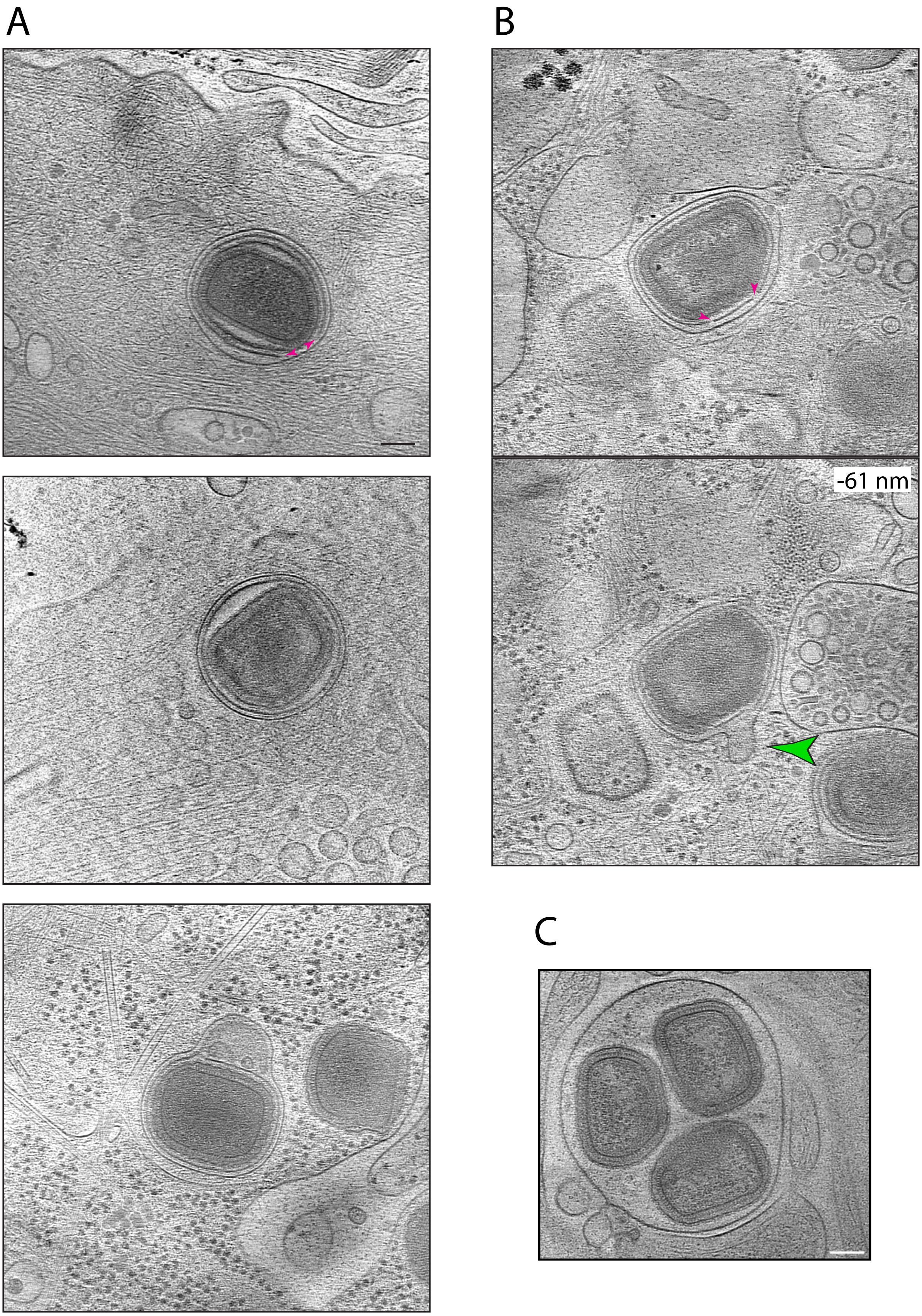


#### Figure S2

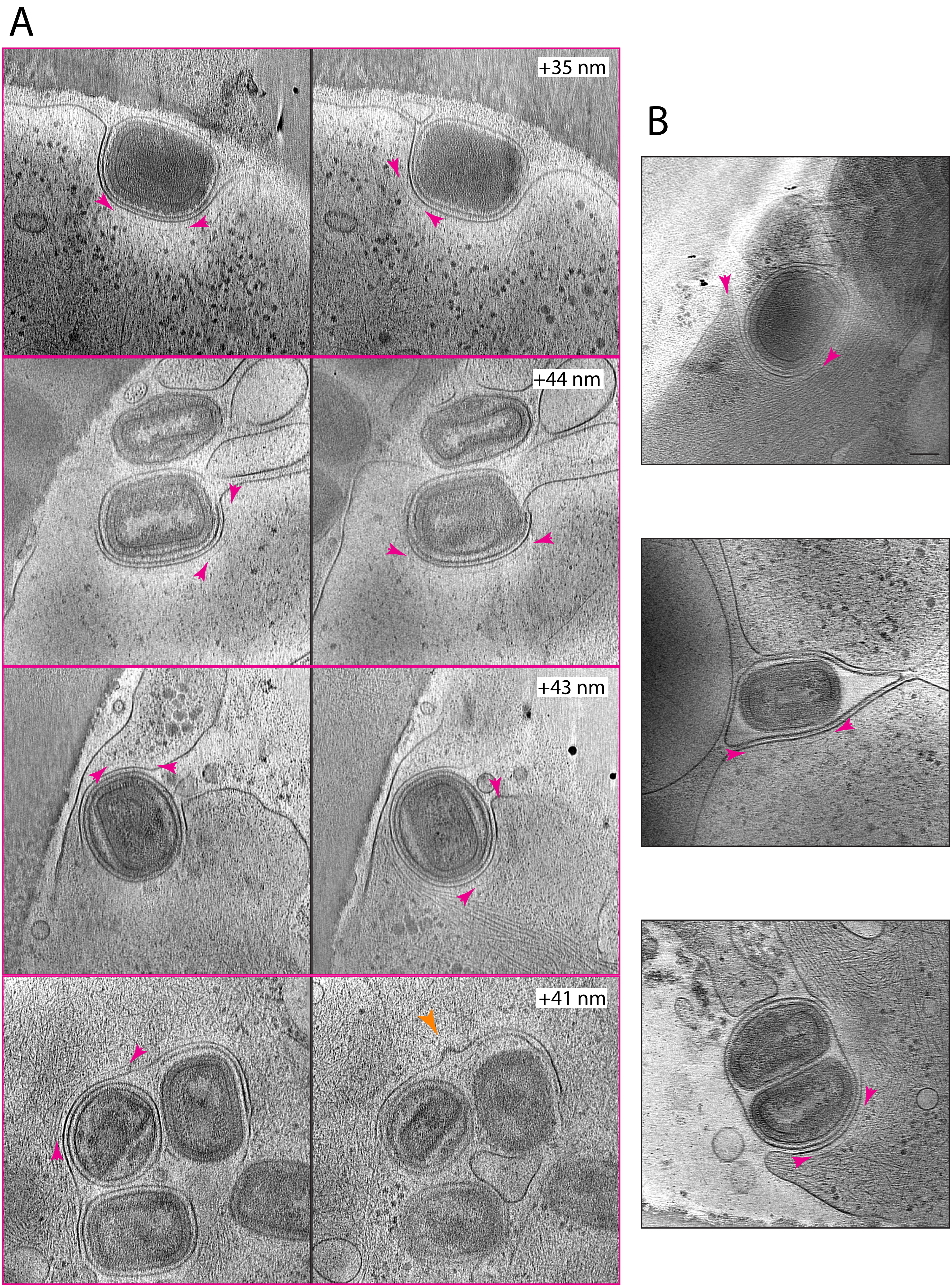


Figure S3

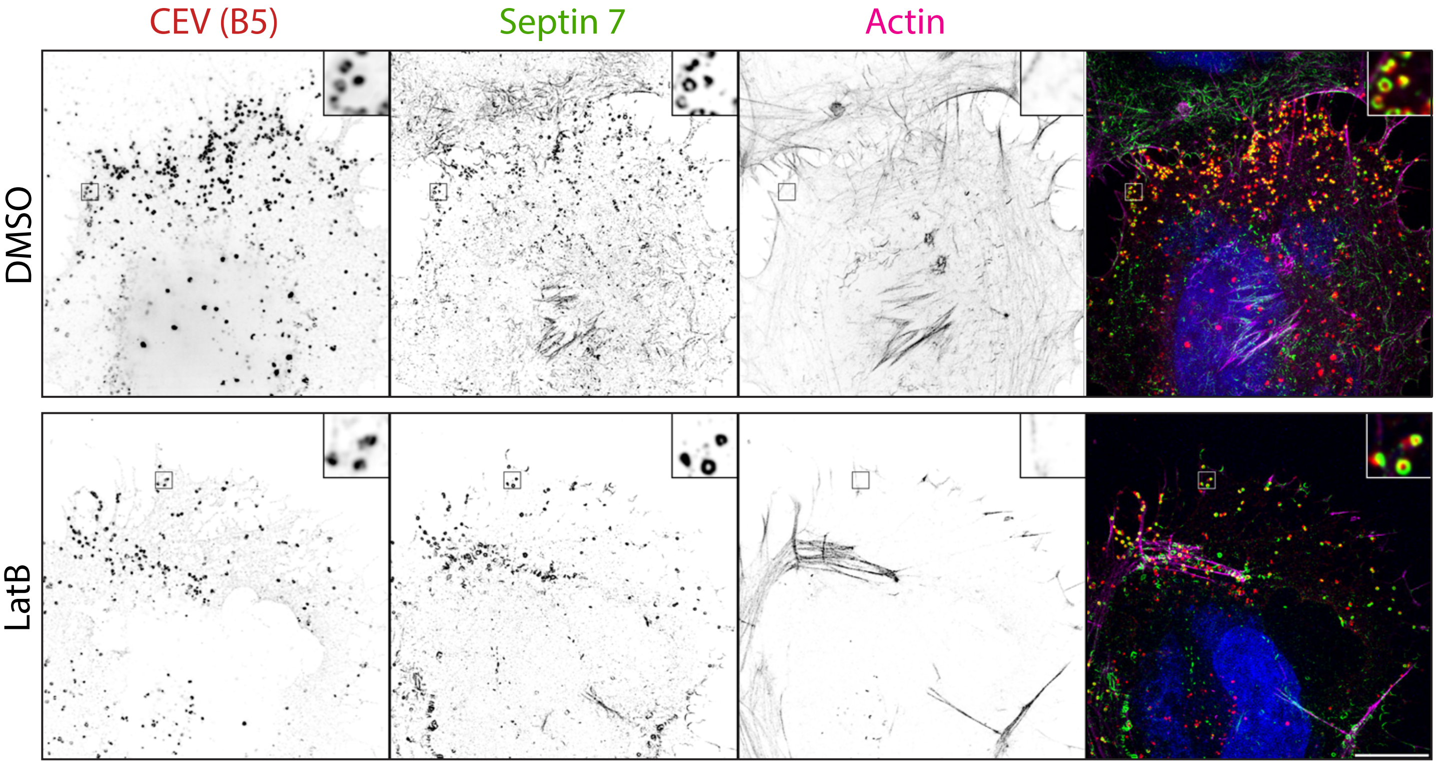


#### Figure S4

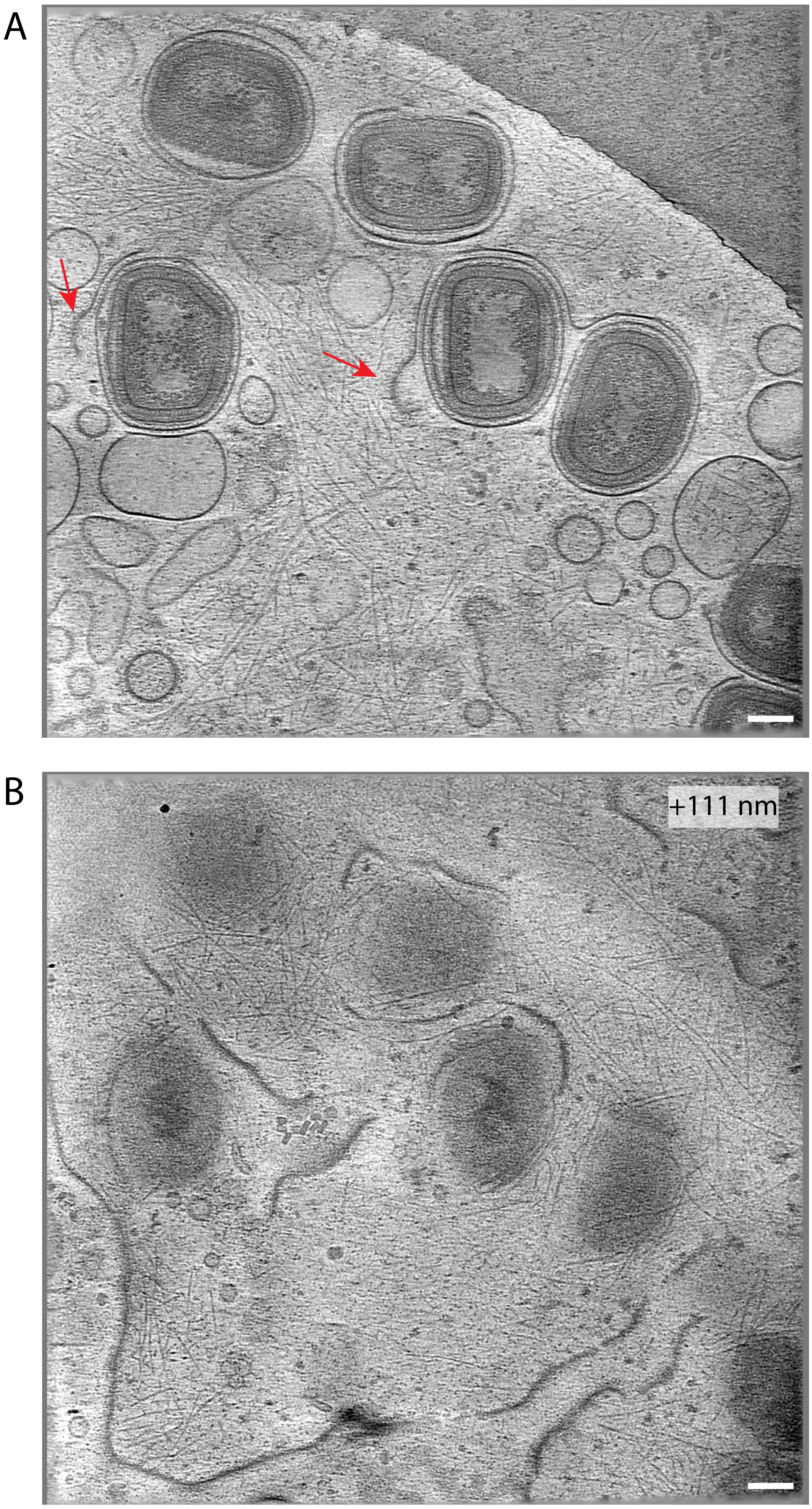


#### Figure S5

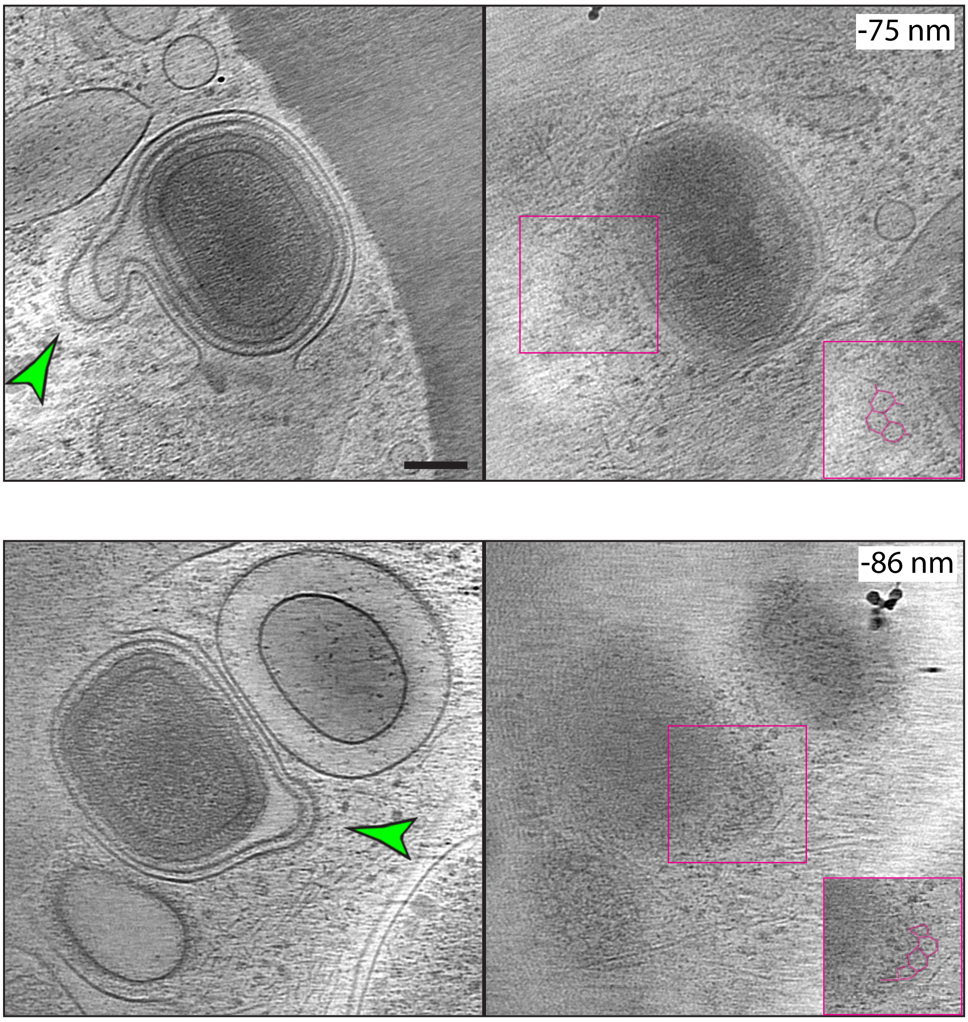


Figure S6
